# nf-core/viralmetagenome: A Novel Pipeline for Untargeted Viral Genome Reconstruction

**DOI:** 10.1101/2025.06.27.661954

**Authors:** Joon Klaps, Philippe Lemey, nf-core community, Liana Eleni Kafetzopoulou

## Abstract

**Motivation:** Eukaryotic viruses present significant challenges for genome reconstruction and variant analysis due to their extensive diversity and potential genome segmentation. While de novo assembly followed by reference database matching and scaffolding is a commonly used approach, the manual execution of this workflow is extremely time-consuming, particularly due to the extensive reference curation required. Here, we address the critical need for an automated, scalable pipeline that can efficiently handle viral metagenomic analysis without manual intervention.

**Results:** We present nf-core/viralmetagenome, a comprehensive viral metagenomic pipeline for untargeted genome reconstruction and variant analysis of eukaryotic DNA and RNA viruses. Viral-metagenome is implemented as a Nextflow workflow that processes short-read metagenomic samples to automatically detect and assemble viral genomes, while also performing variant analysis. The pipeline features automated reference selection, consensus quality control metrics, comprehensive documentation, and seamless integration with containerization technologies, including Docker and Singularity. We demonstrate the utility and accuracy of our approach through validation on both simulated and real datasets, showing robust performance across diverse viral families in metagenomic samples.

**Availability:** nf-core/viralmetagenome is freely available at https://github.com/nf-core/viralmetagenome with comprehensive documentation at https://nf-co.re/viralmetagenome

**Supplementary information:** Supplementary data are available at https://github.com/Joon-Klaps/nf-core-viralmetagenome-manuscript online.

## 1 Introduction

Reconstructing viral genomes from metagenomic sequencing data presents considerable computational challenges, particularly for viruses that exhibit extensive genetic diversity even within a single host [1, 2, 3]. This diversity is further compounded by the prevalence of segmented genomes in viral families like influenza, rotavirus, and bunyaviruses, where individual segments can evolve under distinct selective pressures and reassort, contributing to a complex landscape for genome reconstruction. While pipelines are often designed for a specific virus and their subtypes [4], accurate and complete viral genome reconstruction of samples with unknown references typically requires manual curation of contigs and reference matching [5, 6, 7]. This manual curation process is time-consuming, making it impractical for large-scale metagenome studies or rapid response scenarios that involve emerging viral outbreaks of unknown origin.

To address these limitations, we developed nf-core/viralmetagenome, a comprehensive pipeline specifically designed for untargeted viral genome reconstruction. The pipeline is developed using Nextflow [8] within the nf-core framework [9], ensuring reproducibility through containerization with Docker [10] and Singularity [11], and enabling portability across computational platforms such as local desktops, high-performance clusters and cloud environments.

## 2 Pipeline Description

Nf-core/viralmetagenome implements an automated workflow that performs *de novo* assembly, reference matching through sequence clustering, and iterative refinement by read mapping and consensus calling to reconstruct viral genomes without prior knowledge of the target sequences. The pipeline consists of five major analytical stages: read preprocessing, read metagenomic diversity assessment, contig assembly and scaffolding, iterative consensus refinement with variant analysis, and consensus quality control (Figure 1). The details of the computational tools available at each pipeline stage are described in Supplementary Table 1. While this manuscript highlights key differences between particular tools, the pipeline offers multiple options to accommodate established user workflows and preferences. The complete source code repository is available at https://github.com/nf-core/viralmetagenome.

**Figure 1:**
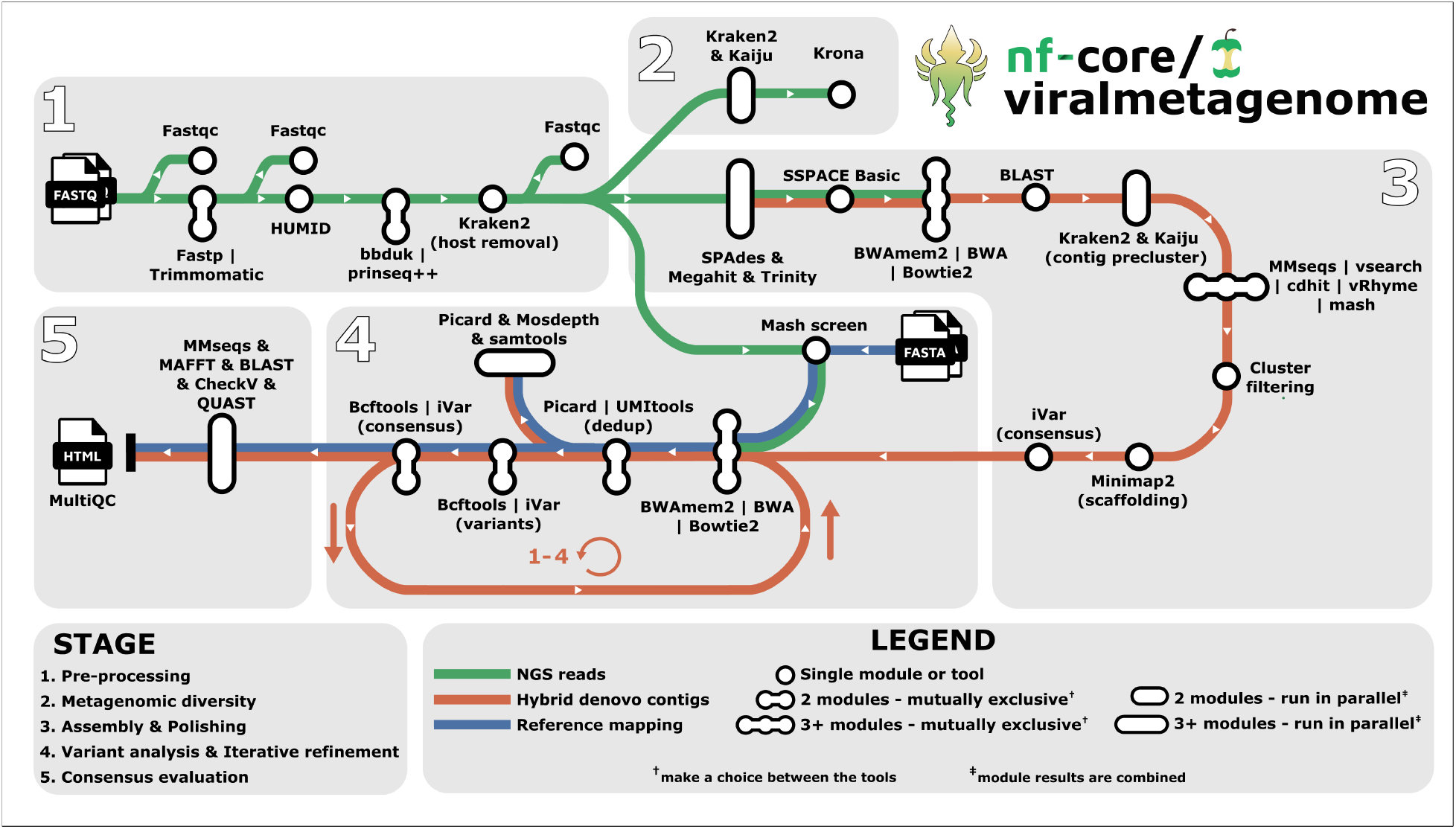
Visual overview of the nf-core/viralmetagenome pipeline for untargeted viral genome reconstruction. nf-core/viralmetagenome processes short-read FASTQ files through optionally read pre-processing (adapter removal, quality filtering, host removal), metagenomic diversity assessment, *de novo* assembly with multiple assemblers, scaffolding with automated reference identification and contig taxonomy-guided clustering, and iterative consensus refinement through read mapping and variant calling. Quality control metrics, assembly statistics, and coverage data are integrated into interactive MultiQC reports and standardised overview tables for downstream analysis.

## 3 Implementation

Nf-core/viralmetagenome only requires nextflow and a container management system (Docker, Singularity, or Conda). The pipeline can be executed with this minimal setup:

~~~
 nextflow run nf-core/viralmetagenome \
 -profile docker \
 --input samplesheet.csv \
 --output results
~~~

Input data is provided through a sample sheet in CSV, TSV, YAML, or JSON format containing sample names and paths to short-read FASTQ files. The pipeline supports both single-end and paired-end sequencing metagenomic data, and offers optional support for Unique Molecular Identifiers (UMIs) as well as optional merging of sequencing runs.

### 3.1 Read preprocessing

The read preprocessing module performs quality control and filtering of raw sequencing reads. Initial quality assessment is conducted using FastQC before and after each processing step to monitor data quality throughout the workflow. Adapter trimming and read processing are performed using either Fastp [12] (default) or Trimmomatic [13]. Fastp is overall faster and has automated adapter detection and trimming [12]. For libraries prepared with UMIs, deduplication is implemented using HUMID [14] and with UMI-tools [15] once reads are mapped to a reference. When multiple sequencing runs from the same sample need to be combined, this can be done after adapter trimming and read-level deduplication by specifying a group identifier in the input samplesheet to indicate which sequencing runs belong to the same biological sample. Complexity filtering, implemented through BBduk [16] or prinseq++ [17], removes low-complexity sequences containing repetitive elements that can lead to spurious alignments or misclassifications during downstream analysis. Host and contamination removal is performed using Kraken2 [18] against a user-specified host genome database. The default database contains a subset of the human genome. However, users are encouraged to employ more comprehensive databases, including complete host genome and transcriptome (human and otherwise), common sequencer contaminants, and bacterial genomes, to ensure thorough decontamination [19].

### 3.2 Metagenomic diversity assessment

Taxonomic classification of preprocessed reads is performed using two complementary approaches - Kaiju [20] and Kraken2 [18] - to maximise detection sensitivity across diverse viral families. Results from both classifiers are visualised using Krona [21].

### 3.3 *De novo* assembly and clustering

The assembly workflow implements a multi-assembler approach followed by clustering and scaffolding procedures. *De novo* assembly is performed using one or multiple assemblers: SPAdes [3] (configured for RNAviral mode by default), MEGAHIT [22], and Trinity [23]. This multi-assembler strategy capitalises on the distinct algorithmic strengths of each tool to maximise genome recovery across diverse viral families and variable read depths. Assembled contigs can be subjected to an optional extension step using SSPACE Basic [24].

Reference identification is conducted through BLASTn [25] against a comprehensive reference sequence pool, with the default being the latest clustered Reference Viral Database [26]. To facilitate identification of related genomic segments and appropriate reference sequences for contig scaffolding, the top five BLAST hits for each contig are retained and incorporated into the subsequent taxonomy-guided clustering step.

Taxonomy-guided clustering employs a two-stage process to cluster related contigs. Initial preclustering uses taxonomic assignments from both Kraken2 [18] and Kaiju [20]. For more efficient targeted analyses, the user can opt to focus on specific taxonomic clades. Subsequent nucleotide similarity clustering is performed using one of six available algorithms: CD-HIT-EST [27], VSEARCH [28], MMseqs2 [29], vRhyme [30], or Mash [31] with network-based community detection using the Leiden algorithm [32] or through single linkage. All tools are valid options, though performance may vary depending on the dataset; for comprehensive benchmarking, we refer to Zielezinski et al. [33] and Steinegger and Söding [29].

As an optional filtering step of contig clusters, after assembly and extension, reads can be mapped to all contigs using BWAmem2 [34] (default), BWA [35], or Bowtie2 [36]. Clusters are filtered based on the cumulated percentage of reads mapped to the contigs of a cluster. By filtering clusters, low-coverage assemblies can be identified that likely represent assembly artefacts.

For the final scaffolding step, all cluster members are mapped to the cluster representative or centroid using Minimap2 [37], followed by consensus calling with iVar [38] to generate reference-assisted assemblies. Regions with zero coverage depth can optionally be represented by the reference genome to produce a more complete scaffold genome for consensus calling.

### 3.4 Iterative consensus refinement and variant calling

The consensus and variant calling module supports two distinct pathways: external reference-based analysis and scaffold refinement. In external reference-based analysis, users can provide reference genomes through the argument `–mapping_constraints`, which allows specifying a separate reference genome or reference set for each sample. When multiple genomes are provided for a single sample, the genome with the highest similarity is used as a reference using Mash [31].

Within the scaffold refinement, the pipeline can perform up to 4 cycles of iterative improvement (default 2) of the scaffolded *de novo* assembled contigs. Each iteration maps reads back to the current consensus using BWAmem2 [34], BWA [35], or Bowtie2 [36], followed by variant calling and consensus generation with BCFtools [39] or iVar [38]. Benchmarking by Bassano et al. [40] showed that BCFtools outperformed iVar in precision and recall. iVar detects more low-frequency variants, resulting in an increased false positive rate but decreases the number of identified false negatives [40]. Users are recommended to consider prioritising sensitivity or specificity when selecting the variant caller.

Optional deduplication can be performed using Picard or when UMI’s are available with UMI-tools [15]. Mapping statistics are generated using samtools (flagstat, idxstats, stats) [39], Picard CollectMultipleMetrics [41], and coverage statistics with mosdepth [42].

### 3.5 Consensus Quality Control

For consensus quality control, nf-core/viralmetagenome employs a multitude of tools. CheckV [43] provides quantitative estimates of genome completeness and contamination levels. Similarity searches are performed using BLASTn [25] against the reference sequence pool and MMseqs2 [29] against comprehensive annotation databases such as Virosaurus [44]. These analyses enable species identification, genomic segment classification, host association determination, and extraction of additional metadata embedded within the reference databases.

The consensus refinement progression can be evaluated through sequence alignment with MAFFT [45], which compares final consensus genomes against *de novo* contigs, intermediate consensus sequences from iterative refinement cycles, and scaffolding references. All tool metrics are compiled into an interactive MultiQC report [46]. Additionally, key metrics are extracted from the MultiQC report and compiled into standalone overview tables to facilitate downstream analysis across all processed samples.

## 4 Applications

To assess the performance of nf-core/viralmetagenome under challenging scenarios, we simulated coinfection scenarios by mixing paired-end reads from public HIV-1 genomes with varying diversity (≤ 80-99% similarity), resulting in 13 samples (See supplementary table 2). Nf-core/viralmetagenome successfully identified coinfections in all mixed samples when genetic similarity was low to moderate (96.7% ANI). We investigated how reference genome selection during scaffolding affects the accuracy of final consensus sequences. When scaffolding references exhibit high similarity (≥ 96%) to the target genome, their influence on the final consensus is minimal. However, more divergent references can introduce substantial errors, with mismatches between final consensus genomes increasing up to 187 nucleotides compared to highly similar references (Figure 2). These findings underscore the critical importance of appropriate reference selection for scaffolding and sequence alignment procedures. Nf-core/viralmetagenome addresses this challenge by automatically selecting the most appropriate reference genome based on sequence similarity; however, when no suitable genome is found in the supplied database, the contigs are scaffolded to a *de novo* contig (specifically, the cluster representative), ensuring that the final consensus genomes are as accurate as possible.

**Figure 2:**
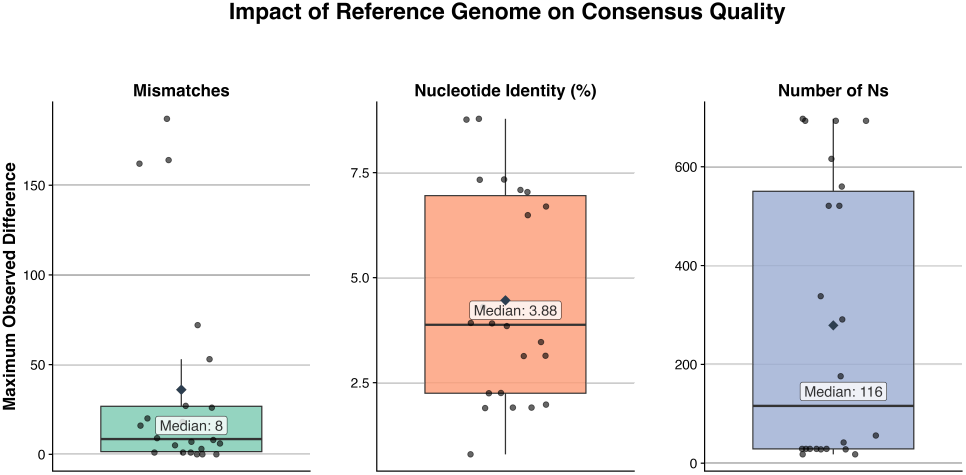
Boxplot of maximum observed differences between consensus sequences generated with different reference genomes during scaffolding. Mean highlighted by a diamond.

To validate nf-core/viralmetagenome’s performance on real-world datasets, we applied the pipeline to publicly available metagenomic samples spanning viral pathogens for both humans and plants. Here, the pipeline successfully generated high-quality or near-complete genomes for all species across different viral families, including both segmented viruses (Lassa virus, Orthonairovirus, Tomato spotted wilt tospovirus) and non-segmented viruses (SARS-CoV-2, West Nile virus, Potato virus Y, Youcai mosaic virus, and Monkeypox virus).

The processing of 28 human viral samples (supplementary methods) required 412 CPU hours and a maximum of 79GB RAM on an HPC system, excluding taxonomic classification steps. The automated reference-contig clustering strategy offers substantial improvements over manual curation approaches by reducing processing time while preserving reconstruction accuracy. Pipeline performance correlates strongly with reference database quality and comprehensiveness—higher-quality, more similar reference genomes produce more complete and accurate consensus sequences. These results emphasize the critical importance of maintaining up-to-date, comprehensive databases such as the Reference Viral Database [26] and Virosaurus [44]. The pipeline uses the Virosaurus-vertebrate database by default for contig annotation; users analysing plant pathogens should consider switching to the Virosaurus-plant database for more accurate results. Since nf-core/viralmetagenome is primarily designed for eukaryotic viruses, bacteriophage analysis requires different approaches and users are encouraged to explore pipelines targeting phages such as VIRify [47], VIBRANT [48], VirSorter2 [49].

## 5 Conclusion

nf-core/viralmetagenome addresses a critical need in viral genomics by providing an automated, scalable solution for untargeted viral genome reconstruction. The pipeline successfully automates the traditionally time-consuming and manual execution process of viral genome assembly from short-read metagenomic data through its integrated workflow of *de novo* contig assembly, automated reference selection, clustering algorithms, and iterative refinement strategies.

Our validation demonstrates the pipeline’s broad applicability across diverse eukaryotic viral families, achieving high-quality genome reconstruction while ensuring reproducibility and ease of deployment across different computational environments.

As viral surveillance and outbreak response increasingly rely on metagenomic sequencing, automated pipelines like nf-core/viralmetagenome will be essential for the timely identification of pathogen strains. The pipeline represents a significant step forward in making viral genome reconstruction accessible to researchers without requiring extensive bioinformatics expertise, facilitating broader adoption of metagenomic approaches in viral research and public health applications.

## Acknowledgments

J.K, P.L. and L.E.K acknowledge support from the Research Foundation - Flanders (Fonds voor Wetenschappelijk Onderzoek – Vlaanderen, G005323N and G051322N, 1SH2V24N, 12×9222N).

## Author Contributions

J.K. designed and implemented the pipeline, performed validation analyses, and wrote the manuscript. P.L. and L.E.K. supervised the project and provided critical feedback. The nf-core community contributed to maintaining the pipeline. All authors reviewed and approved the final manuscript.

## Data Availability

The nf-core/viralmetagenome pipeline is freely available at https://github.com/nf-core/viralmetagenome. The raw data and analysis code is available on https://github.com/Joon-Klaps/nf-core-viralmetagenome-manuscript/.

