## Supplementary Table for "nf-core/viralmetagenome: A Novel Pipeline for Untargeted Viral Genome Reconstruction"

### Supplementary Methods & Tables - nf-core/viralmetagenome: A Novel Pipeline for Untargeted Viral Genome Reconstruction

[2]A full list of contributors can be found at https://nf-co.re/community.

June 27, 2025

### Supplementary Methods

#### S1. Influence of Scaffold Reference on Consensus Genome

We simulated mixed HIV-1 samples containing paired-end read data based on four genomes (MN090277.1, MN090188.1, MN090240.1, and MZ766668.1) spanning genetic diversity from 80–99% average nucleotide identity (ANI). Thirteen sample configurations were created with varying abundance ratios to model both single-strain infections (100% abundance) and coinfections with different abundance ratios (75:25, 50:50, 25:75) to reflect coinfections ( Supplementary table 1). Reads were generated using InSilisocoSeq v2.0.1 [1] using 'kde' mode and 'lognormal' abundance for a 'MiSeq' model.

Sample mixtures were run with nf-core/viralmetagenome using default settings unless specified otherwise. Reference sequences used to compare the influence of the reference during scaffolding are the same sequences used for read generation, in addition to the databases, Virosaurus [2] and the Reference Viral Database [3]. Visualisation was done in R [4] using tidyverse [5].

#### S2. Evaluation of nf-core/viralmetagenome on public data

A public metagenomic dataset was constructed by selecting a randomized subset of 2,000 records from the NCBI Virus database, targeting the following viruses: *Orthonairovirus haemorrhagiae*, *Mammarenavirus lassaense*, *Zika virus*, *West Nile virus*, *Monkeypox virus*, *Influenza A virus*, *Severe acute respiratory syndrome coronavirus 2*, and *Human respiratory syncytial virus A*. Records were filtered to retain only those with associated SRA data, sequenced on the Illumina platform, and with a LibraryStrategy other than 'Amplicon'. From the resulting pool, 28 samples were randomly selected and downloaded using nf-core/fetchngs. Samples were then run with nf-core/viralmetagenome using `-cluster_method 'mmseqs-cluster'` due to sequence size limitations of cdhit, `-keep_unclassified false` and `-skip_read_classification` to speed up the analysis.

For plant pathogens, two samples were randomly selected from the SRA after querying for the following viruses: Tobacco mosaic virus, Tomato spotted wilt virus, Cucumber mosaic virus, and Potato virus Y. These were also downloaded using nf-core/fetchngs and processed with viralmetagenome using the same parameters, with the addition of the Virosaurus plant virus database.

### Supplementary Tables

Table 1: Overview of computational tools and methods used in the nf-core/viralmetagenome pipeline. The pipeline integrates multiple bioinformatics tools across different stages of viral metagenomic analysis, from preprocessing to consensus calling. Each tool was selected for its specific capabilities and performance characteristics relevant to viral sequence analysis.

| Stage | Step | Tool | Explanation | Reference |
|---|---|---|---|---|
| Pre-processing | Read trimming | fastp | A high-performance preprocessing solution built in C++ that consolidates multiple quality control operations into a single workflow. Fastp delivers enhanced processing speed compared to traditional alternatives, like Trimmomatic, while handling adapter removal, quality assessment, base correction, read and filtering. Fastp also features automated adapter detection capabilities for various Illumina sequencing protocols. | [6] |
| | | Trimmomatic | A comprehensive preprocessing solution tailored for Illumina sequencing platforms with specialised paired-end read handling capabilities. Provides multiple pre-processing options, including adapter removal, quality-based trimming using sliding window approaches, and length-based filtering. | [7] |
| | Complexity filtering | Bbduk | A high-performance, multi-threaded tool designed to combine various data-quality-related operations, including trimming, filtering, and masking, into a single pass. It is particularly effective for removing host and other contaminant DNA using k-mer matching strategies, which is crucial for accurate metagenomic analysis. | [8] |
| | | Prinseq++ | A C++ implementation that significantly improves upon its predecessor, prinseq-lite.pl, in terms of computational efficiency. It offers extensive quality control features, including filtering by length, GC content, quality scores, N content, and sequence complexity (entropy/DUST score). | [9] |
| Metagenomic diversity | Taxonomy classification | Kraken2 | An advanced taxonomic classification system with an improved k-mer-based approach through optimised memory usage and enhanced processing speed. Incorporates protein-level search capabilities for improved detection of divergent viral sequences, with a library of pre-built RefSeq indexes. | [10] |

*Continued on next page*

*Table 1 continued from previous page*

| Stage | Step | Tool | Explanation | Reference |
|---|---|---|---|---|
| | | Kaiju | A protein-level taxonomic classifier that employs exact matching algorithms on translated sequences using the Burrows-Wheeler transform. It can optionally allow amino acid substitutions and tends to have higher sensitivity and precision compared to k-mer-based classifiers (Kraken2). Kaiju is particularly effective for organisms with limited reference representation. | [11] |
| Assembly & polishing | Assembly | SPAdes | A de Bruijn graph-based assembler designed to overcome challenges in complex sequencing data, such as non-uniform coverage (using multisized de Bruijn graphs) and variable insert sizes. The Spades suite contains multiple specialised modes rnaSPAdes, coronaSPAdes, metaSPAdes for extra flexibility. | [12] |
| | | MEGAHIT | A succinct de Bruijn graph-based assembler specifically designed for large and complex metagenomic datasets. MEGAHIT has demonstrated the ability to generate larger assemblies with longer contig N50 and average contig length. Its relatively low memory requirements make it well-suited for handling large datasets. | [13] |
| | | Trinity | A powerful modular transcriptome reconstruction tool, capable of fully reconstructing a large fraction of transcripts, including alternative splice isoforms and transcripts from recently duplicated genes, even in the absence of a reference genome. It partitions sequence data into individual de Bruijn graphs, processing each independently to extract full-length isoforms and resolve paralogous genes. | [14] |
| | Read alignment | BWAmem2 | An optimized version of the BWA-MEM algorithm enhanced with Intel-specific acceleration feature, providing approximately a twofold speedup in alignment throughput over BWA-MEM while ensuring identical SAM output. It is highly efficient for aligning short reads to large reference genomes. | [15] |

*Continued on next page*

*Table 1 continued from previous page*

| Stage | Step | Tool | Explanation | Reference |
|---|---|---|---|---|
| | | BWA | A foundational and widely used Burrows-Wheeler Aligner (BWA-MEM variant) that efficiently maps short reads and long sequences to large reference genomes. It supports long gaps and dynamically adjusts error rates based on sequence length, making it robust to sequencing errors. | [16] |
| | | Bowtie2 | A fast and memory-efficient alignment tool for aligning sequencing reads to long reference sequences, such as mammalian genomes. Bowtie2 supports gapped, local, and paired-end alignment modes, combining high speed, sensitivity, and accuracy. | [17] |
| | Clustering | CD-HIT-EST | A highly efficient and widely used program for clustering large sets of protein or nucleotide sequences. Implementation constraints limit processing to genomes under 10Mbp with minimum 0.8. The program employs a greedy incremental algorithm. | [18] |
| | | MMSeqs | A powerful software suite for fast and sensitive deep clustering and searching of large protein sequence sets. It is significantly faster than other tools like BLASTclust and can cluster large databases down to low sequence identity thresholds. MMseqs offers a variety of clustering modes for a lot of flexibility. | [19] |
| | | VSEARCH | An open-source, versatile alternative for USEARCH. VSEARCH supports processing very large datasets, limited primarily by available memory. | [20] |
| | | Mash | A non-alignment based clustering technique that applies MinHash algorithms for rapid genome and metagenome distance estimation. It compresses large sequences into small, representative sketches, enabling ultra-fast estimations of global mutation distances. Mash creates compact sequence representations enabling ultra-fast similarity assessments and large-scale clustering operations, though accuracy decreases below 95% genome similarity. | [21] |

*Continued on next page*

*Table 1 continued from previous page*

| Stage | Step | Tool | Explanation | Reference |
|---|---|---|---|---|
| | | vRhyme | A machine learning-based binning solution specifically developed for viral genome recovery from metagenomic datasets. VRhyme addresses the unique challenges of viral sequences, such as the lack of universal marker genes. Addresses unique viral sequence characteristics through supervised learning approaches combined with coverage-based analysis across multiple samples. Generates high-quality viral bins with minimal computational overhead, though it may produce empty results when insufficient viral evidence is detected. | [22] |
| Variant & consensus calling | Variant & consensus calling | iVar | A computational package specifically designed for viral amplicon-based sequencing, integrating functions like consensus and variant calling (including iSNVs and insertions/deletions). It is a key component of best-practice pipelines for reconstructing consensus genomes from viral sequencing data. | [23] |
| | | BCFtools | A robust suite of utilities for manipulating variant calls in VCF and BCF formats. Provides extensive functionality for variant calling using multiallelic models, data filtering, file merging, and consensus sequence generation through variant application. | [24] |

Table 2: Overview of simulated HIV-1 sample mixtures with the used reference genome and metadata.

| **Sample name** | **Reference** | **Added genome** | **Similarity to reference (%)** | **Reads (M)** | **Read % MN090277** | **Read % added genome** |
|---|---|---|---|---|---|---|
| sample_1 | MN090277.1 | MN090188.1 | 98.7 | 4 | 75 | 25 |
| sample_2 | MN090277.1 | MN090188.1 | 98.7 | 4 | 50 | 50 |
| sample_3 | MN090277.1 | MN090188.1 | 98.7 | 4 | 25 | 75 |
| sample_4 | MN090277.1 | MN090240.1 | 96.7 | 4 | 75 | 25 |
| sample_5 | MN090277.1 | MN090240.1 | 96.7 | 4 | 50 | 50 |
| sample_6 | MN090277.1 | MN090240.1 | 96.7 | 4 | 25 | 75 |
| sample_7 | MN090277.1 | MZ766668.1 | 83.5 | 4 | 75 | 25 |
| sample_8 | MN090277.1 | MZ766668.1 | 83.5 | 4 | 50 | 50 |
| sample_9 | MN090277.1 | MZ766668.1 | 83.5 | 4 | 25 | 75 |
| sample_10 | MN090277.1 | MN090277.1 | 100.0 | 1 | 100 | 0 |
| sample_11 | MN090277.1 | MN090188.1 | 98.7 | 1 | 0 | 100 |
| sample_12 | MN090277.1 | MN090240.1 | 96.7 | 1 | 0 | 100 |
| sample_13 | MN090277.1 | MZ766668.1 | 83.5 | 1 | 0 | 100 |

### References

[1] Hadrien Gourlé, Oskar Karlsson-Lindsjö, Juliette Hayer, and Erik Bongcam-Rudloff. Simulating illumina metagenomic data with InSilicoSeq. 35:521–522.

[2] Anne Gleizes, Florian Laubscher, Nicolas Guex, Christian Iseli, Thomas Junier, Samuel Cordey, Jacques Fellay, Ioannis Xenarios, Laurent Kaiser, and Philippe Le Mercier. Virosaurus a reference to explore and capture virus genetic diversity. 12.

[3] Norman Goodacre, Aisha Aljanahi, Subhiksha Nandakumar, Mike Mikailov, and Arifa S Khan. A reference viral database (RVDB) to enhance bioinformatics analysis of high-throughput sequencing for novel virus detection. 3.

[4] R Core Team. R: A language and environment for statistical computing.

[5] Hadley Wickham, Mara Averick, Jennifer Bryan, Winston Chang, Lucy D'agostino McGowan, Romain François, Garrett Grolemund, Alex Hayes, Lionel Henry, Jim Hester, Max Kuhn, Thomas Lin Pedersen, Evan Miller, Stephan Milton Bache, Kirill Müller, Jeroen Ooms, David Robinson, Dana Paige Seidel, Vitalie Spinu, Kohske Takahashi, Davis Vaughan, Claus Wilke, Kara Woo, and Hiroaki Yutani. Welcome to the tidyverse.

[6] Shifu Chen, Yanqing Zhou, Yaru Chen, and Jia Gu. fastp: an ultra-fast all-in-one FASTQ preprocessor. 34:i884–i890.

[7] Anthony M Bolger, Marc Lohse, and Bjoern Usadel. Trimmomatic: a flexible trimmer for illumina sequence data. 30:2114–2120.

[8] Brian Bushnell. BBMap.

[9] Vito Adrian Cantu, Jeffrey Sadural, and Robert Edwards. PRINSEQ++, a multi-threaded tool for fast and efficient quality control and preprocessing of sequencing datasets.

[10] Derrick E Wood, Jennifer Lu, and Ben Langmead. Improved metagenomic analysis with kraken 2. 20:257.

[11] Peter Menzel, Kim Lee Ng, and Anders Krogh. Fast and sensitive taxonomic classification for metagenomics with kaiju. 7:11257.

[12] Dmitry Meleshko, Iman Hajirasouliha, and Anton Korobeynikov. coronaSPAdes: from biosynthetic gene clusters to RNA viral assemblies.

[13] Dinghua Li, Ruibang Luo, Chi-Man Liu, Chi-Ming Leung, Hing-Fung Ting, Kunihiko Sadakane, Hiroshi Yamashita, and Tak-Wah Lam. MEGAHIT v1.0: A fast and scalable metagenome assembler driven by advanced methodologies and community practices. 102:3–11.

[14] Manfred G Grabherr, Brian J Haas, Moran Yassour, Joshua Z Levin, Dawn A Thompson, Ido Amit, Xian Adiconis, Lin Fan, Raktima Raychowdhury, Qiandong Zeng, Zehua Chen, Evan Mauceli, Nir Hacohen, Andreas Gnirke, Nicholas Rhind, Federica di Palma, Bruce W Birren, Chad Nusbaum, Kerstin Lindblad-Toh, Nir Friedman, and Aviv Regev. Full-length transcriptome assembly from RNA-seq data without a reference genome. 29:644–652.

[15] Md Vasimuddin, Sanchit Misra, Heng Li, and Srinivas Aluru. Efficient architecture-aware acceleration of BWA-MEM for multicore systems. In *2019 IEEE International Parallel and Distributed Processing Symposium (IPDPS)*, pages 314–324. IEEE.

[16] Heng Li. Aligning sequence reads, clone sequences and assembly contigs with BWA-MEM.

[17] Ben Langmead, Christopher Wilks, Valentin Antonescu, and Rone Charles. Scaling read aligners to hundreds of threads on general-purpose processors. 35:421–432.

[18] Weizhong Li and Adam Godzik. Cd-hit: a fast program for clustering and comparing large sets of protein or nucleotide sequences. 22:1658–1659.

[19] Martin Steinegger and Johannes Söding. MMseqs2 enables sensitive protein sequence searching for the analysis of massive data sets. 35:1026–1028.

[20] Torbjørn Rognes, Tomáš Flouri, Ben Nichols, Christopher Quince, and Frédéric Mahé. VSEARCH: a versatile open source tool for metagenomics. 4:e2584.

[21] Brian D Ondov, Gabriel J Starrett, Anna Sappington, Aleksandra Kostic, Sergey Koren, Christopher B Buck, and Adam M Phillippy. Mash screen: high-throughput sequence containment estimation for genome discovery. 20:232.

[22] Kristopher Kieft, Alyssa Adams, Rauf Salamzade, Lindsay Kalan, and Karthik Anantharaman. vRhyme enables binning of viral genomes from metagenomes. 50:e83.

[23] Nathan D Grubaugh, Karthik Gangavarapu, Joshua Quick, Nathaniel L Matteson, Jaqueline Goes De Jesus, Bradley J Main, Amanda L Tan, Lauren M Paul, Doug E Brackney, Saran Grewal, Nikos Gurfield, Koen K A Van Rompay, Sharon Isern, Scott F Michael, Lark L Coffey, Nicholas J Loman, and Kristian G Andersen. An amplicon-based sequencing framework for accurately measuring intrahost virus diversity using PrimalSeq and iVar. 20:8.

[24] Petr Danecek, James K Bonfield, Jennifer Liddle, John Marshall, Valeriu Ohan, Martin O Pollard, Andrew Whitwham, Thomas Keane, Shane A McCarthy, Robert M Davies, and Heng Li. Twelve years of SAMtools and BCFtools. 10.
